# Glucose alters the evolutionary response to gentamicin in uropathogenic *Escherichia coli*

**DOI:** 10.1101/2024.11.26.625364

**Authors:** Shalini Choudhary, Jacob A. Smith, Alan McNally, Rebecca J. Hall

## Abstract

Urinary tract infections (UTI) are a major health and economic concern. Uropathogenic *Escherichia coli* (UPEC) are the leading cause of UTI, and antibiotic resistant UPEC are increasingly common. The microenvironment of the urinary tract is metabolically distinct, and there is growing interest in understanding the extent to which metabolism may influence UPEC infection and response to antibiotics, and how this varies between individuals. Diabetes, characterised in part by glycosuria, is a known risk factor for UTI and is associated with more severe infections. The role that glucose plays in driving UPEC evolution is unclear. Here, we found that a pathologically-relevant glucose concentration reduced the efficacy of the antibiotic gentamicin against a UPEC strain. Through experimental evolution, we identified mutations in the RNA polymerase sigma factor *rpoS* associated with long-term glucose exposure. We found that the presence of gentamicin resulted in mutations in genes including *trkH*, encoding a potassium ion uptake system and linked previously to aminoglycoside resistance, and in the autotransporter *hyxB*. Strikingly, these mutations were not present in populations exposed to a combination of both glucose and gentamicin. Together, this suggests that whilst glucose may reduce growth inhibition by gentamicin, it may also influence mutation acquisition, providing new avenues for understanding the evolution and treatment of UPEC-mediated UTI in high-risk individuals.

## Introduction

Urinary tract infections (UTI) are a major cause of morbidity and mortality globally (1; 2). Certain demographics are disproportionately affected, including women and individuals with indwelling catheters or diabetes (3; 4; 5; 6), and recurrent infections are common (3). UTI are treated by a panel of antibiotics depending on the acuteness and severity of infection, and whether it has progressed to the bloodstream causing urosepsis (7; 8). Uropathogenic *Escherichia coli* (UPEC) are the leading cause of UTI, with antibiotic and multidrug resistant UPEC strains increasingly common (7; 9; 10; 11).

Whilst virulence factors are known to contribute to UPEC pathogenicity (2; 12; 13; 14), less is known about the extent to which other genes and pathways influence infection, evolution, and antibiotic resistance. The urinary tract is uniquely composed of metabolites including urea, uric acid, ammonia, and creatinine, thereby creating a distinct metabolic microenvironment to which UPEC are exposed (15; 16). The concentration of glucose in urine can also become elevated (glycosuria, defined as greater than 25 mg/dL (17)) as a result of conditions including diabetes mellitus and gestational diabetes. Genes involved in glucose transport and metabolism have been shown to be important for UPEC colonisation of the urinary tract, including *ptsG* (a component of a glucose-specific phosphotransferase system [PTS]) and *pgi* (catalyses the reversible reaction between glucose 6-phosphate and fructose 6-phosphate) (16; 18; 19; 20). Given the increased risk of UTI amongst this demographic (21), it is important to understand the extent to which glycosuria might influence antibiotic susceptibility or drive the evolution of resistance in UPEC.

Using a clinical UPEC strain, we identified that the presence of glucose prevented a significant reduction in growth by sub-inhibitory concentrations of the aminoglycoside gentamicin, an antibiotic used to treat severe UTI or urosepsis. We then experimentally evolved the UPEC strain in the presence of either a physiologically-relevant concentration of glucose, gentamicin, or both glucose and gentamicin. Analysis of short read sequences of strains from the evolved populations showed that glucose exposure led to mutations in the RNA polymerase sigma factor *rpoS*, and gentamicin to mutations in the autotransporter *hyxB* and the potassium ion uptake system *trkH*. Crucially, no mutations in *trkH* nor *hyxB* were identified in the populations evolved in both glucose and gentamicin. This suggests therefore that metabolism may play a role in the treatment and evolution of UPEC-associated UTI.

## Methods

### Strains and growth conditions

To measure potential effect of glucose on antibiotic sensitivity, *E. coli* USVAST002, originally isolated from a human urine sample (22), was first streaked from a glycerol stock on to a Luria Bertani (LB) agar plate (E & O Laboratories Ltd), and incubated overnight at 37°C. A single colony was used to inoculate 5 mL LB broth (E & O Laboratories Ltd) in a 30 mL universal before overnight incubation at 37°C with agitation. The overnight culture was then diluted 1:1000 in LB. Two times final concentrations of glucose (2 mg/mL) and gentamicin (500 ng/mL, 250 ng/mL, and 100 ng/mL) were added to LB, 50 *µ*L of which was then added to each test well of a 96-well plate in triplicate, followed by 50 *µ*L of the dilute cell suspension to a final volume of 100 *µ*L per well. Glucose- and gentamicin-free conditions were also included. Plates were incubated for 24 hours in a microplate reader (Tecan) at 37°C with continuous double orbital shaking, with absorbance measurements (OD600) taken every 30 minutes in triplicate. Minimum inhibitory concentrations (MIC) of the ancestral and evolved strains were estimated in U-bottom 96-well plates (23). For this, overnight cultures were diluted 1:2000 in Iso-Sensitest broth (ISB) (Thermo Fisher Scientific) and incubated in triplicate in seven different concentrations of gentamicin ranging from 16 *µ*g/mL to 250 ng/mL, including a no-antibiotic control. MICs were recorded following an 18-hour incubation at 37°C, with significance considered if results were two doubling dilutions away from the ancestor.

### Experimental evolution

The ancestral *E. coli* USVAST002 isolate was streaked from a glycerol stock on to an LB agar plate and incubated overnight at 37°C. A single colony was used to inoculate 5 mL LB in a 30 mL universal, with six independent biological replicates per condition: LB no antibiotic (henceforth ‘control’); LB + 2 mg/mL added glucose (‘Glc’); LB + 500 ng/mL gentamicin (‘GEN’); LB + 2 mg/mL glucose + 500 ng/mL gentamicin (‘Glc+GEN’). Microcosms were incubated for 24 hours at 37°C with agitation, before a 1% transfer of cell suspension into fresh media. This 1% transfer was repeated every 24 hours for 21 days. Every seven days, the whole population was centrifuged at 3600 rpm (Thermo Scientific Megafuge 40R TX-1000) for five minutes, resuspended in 1 mL 50% glycerol, and stored at -80°C.

### Genome sequencing and bioinformatics

Sequencing of the ancestor (hybrid) and evolved (short read) strains was performed by MicrobesNG (UK). A single colony of the ancestral strain was sequenced from pure culture. For the six control populations, a single colony was randomly selected. For each of the test populations, the population was first grown to single colonies on an LB agar plate before 16 colonies per population were added directly to 100 *µ*L of 1 *µ*g/mL gentamicin in LB in individual wells of a 96-well plate. The plate was then incubated for 24 hours at 37°C with agitation, the resulting kinetics plotted, and three strains selected for sequencing that best represented the phenotypic spread. The growth assay was then repeated in triplicate for each of the selected strains. The ancestor hybrid assembly was generated using Flye (v2.8.1-b1676) (24), polished using Polypolish (v0.5.0) (25), and annotated with Bakta (v1.8.2) (26). Genes of interest that were assigned a function but not a gene name were confirmed by BLASTn queries. Variants were called against the assembled ancestor using breseq (v0.36.0) (27) with a consensus minimum variant coverage of 10 and a minimum mapping quality of 20. Genes with variants that were present in the control populations were discounted from the test populations. Resistance genes in the ancestral strain were predicted using ABRicate (v0.8) to query the ResFinder database, and sequence type (ST) ascertained using mlst (v2.23.0) (28).

### Statistical analyses

Area under the curve measurements were calculated using numpy.trapz in Python (v3.9.10). Significance testing was conducted using a one-way analysis of variance (ANOVA).

## Results

### Glucose rescues growth in sub-inhibitory gentamicin

USVAST002 (ST73) was selected for its status as a UPEC strain with no predicted functional resistance genes. To assess how the action of gentamicin is affected by moderate glycosuria (29), USVAST002 was grown to 2 mg/mL glucose plus subinhibitory gentamicin (100 ng/mL, 250 ng/mL, or 500 ng/mL). At increasing concentrations of gentamicin there was a corresponding reduction in growth (Fig. 1). In the absence of gentamicin, the addition of glucose increased growth significantly. When USVAST002 was grown in both glucose and gentamicin, there was no significant difference between that condition and the cells grown in glucose alone. This suggests that glucose acts to reduce the effect of gentamicin.

**Fig. 1.**
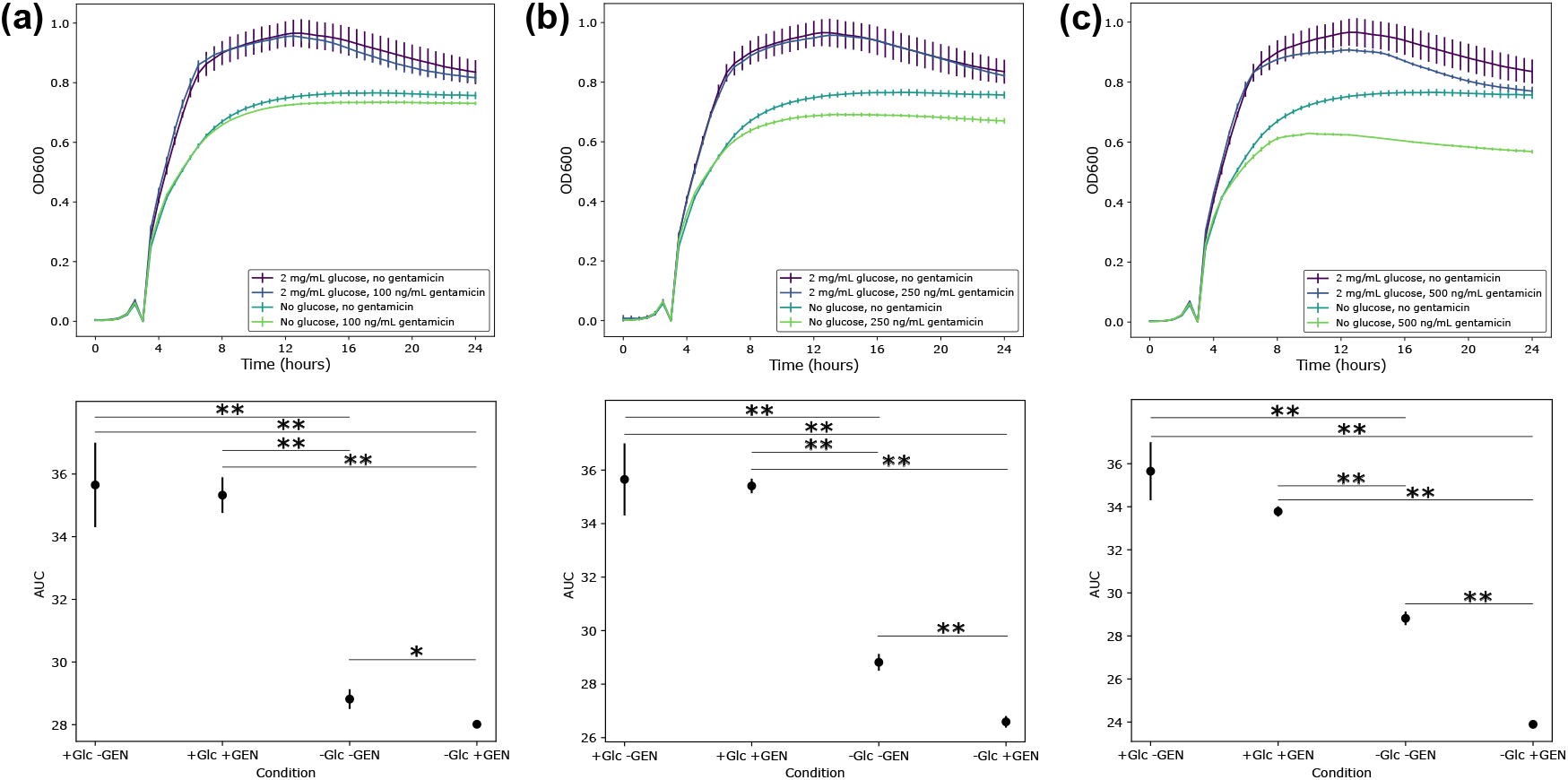
Growth kinetics (top) of USVAST002 and corresponding area under the curve (AUC, bottom) values in 2 mg/mL added glucose plus (a) 100 ng/mL, (b) 250 ng/mL, or (c) 500 ng/mL gentamicin. Measurements in triplicate, error bars standard deviation. * p < 0.05, ** p < 0.01, one-way ANOVA.

### Phenotypic changes following prolonged gentamicin exposure

To assess how glucose might affect the evolution of gentamicin resistance, we performed a 21-day evolution experiment in conditions of 2 mg/mL glucose (Glc), 500 ng/mL gentamicin (GEN), and both glucose and gentamicin (Glc+GEN). We then screened the growth kinetics of three colonies from each of the evolved populations in LB only and in 1 *µ*g/mL gentamicin, as well as performing MIC assays in a gradient of gentamicin. In these conditions, the majority of the strains from the Glc and the Glc+GEN populations had comparably worse growth than the ancestor (Fig. S1, Fig. S2). In contrast, with the exception of strain GEN 4C, most of the strains from the GEN populations grew comparably to the ancestor in LB only, but seven strains grew significantly better in 1 *µ*g/mL gentamicin. GEN 4C also grew more poorly than the ancestor in the presence of antibiotic. No strain had an MIC that was significantly different to the ancestor (Table S1).

### Long-term exposure to glucose causes significant genetic changes

The extent to which glucose may act as a selective pressure is not understood fully. To investigate this, we performed short read sequencing of the three individual colonies from each of the evolved populations and analysed the significant genetic changes. The changes were pooled for each replicate to gain a representative understanding of mutations that were present in the population as a whole (Fig. 2), but are provided at an individual strain level in Table S1. Mutations in the RNA polymerase sigma factor *rpoS* were identified in five of the six replicate populations in both the Glc and the Glc+GEN condition. This included five different non-synonymous single nucleotide polymorphisms (SNPs) that introduced premature stop codons (Glc P1 Q59*, and Glc+GEN P1 Q251*, P3 Q257*, P4 L30*) (Table S1). In the sixth Glc population, there was an intergenic SNP between *nlpD* and *rpoS*, and in the sixth Glc+GEN a 6,506 base pair (bp) deletion spanned the same region. No changes were observed in this region in any of the control or GEN populations, indicating that they may have arisen as a result of glucose exposure.

**Fig. 2.**
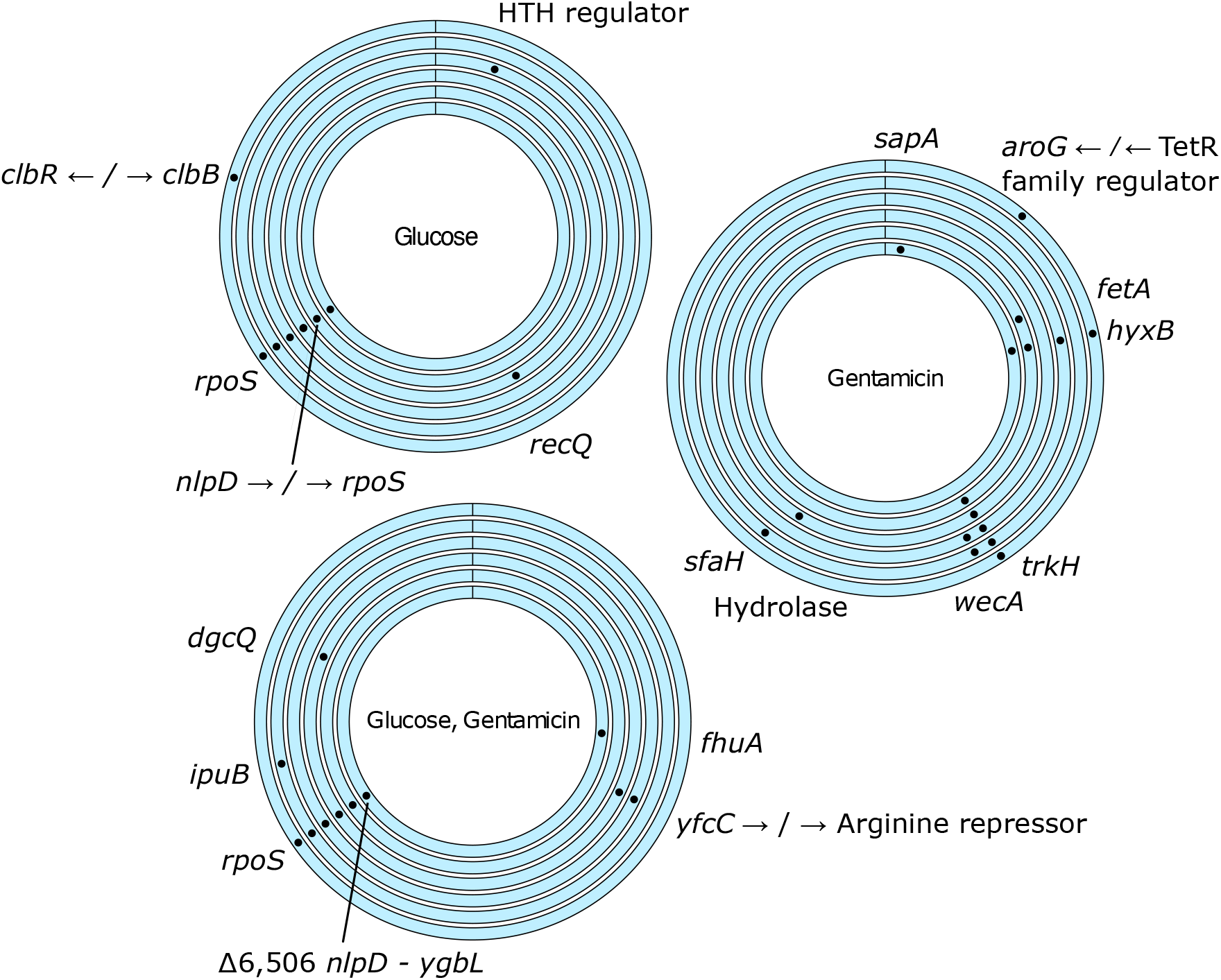
Genetic changes identified in response to glucose, gentamicin, or both glucose and gentamicin. Each ring represents pooled changes from three individual colonies from each replicate population. Changes also identified in any control population are not presented here.

### Glucose alters the evolutionary response to gentamicin

We then analysed the populations evolved in the presence of gentamicin to identify any significant genetic changes arising as a result of antibiotic exposure. We identified SNPs in *trkH*, encoding a potassium ion uptake system, in five of the six GEN populations (Fig. 2). There were four different non-synonymous mutations identified across the replicates (I15L, L185Q, G25R, P151S), with one GEN population carrying at least two different mutations (GEN 1A I15L, GEN 1B L185Q) (Table S1). We also found mutations in *hyxB*, an autotransporter, in four of the six populations, and two in *wecA* (involved in enterobacterial common antigen and O-antigen biosynthesis). Strains with three of these *trkH* mutations had significantly better growth in the presence of 1 *µ*g/mL gentamicin (Fig. 3). In contrast, none of the strains carrying mutations in *hyxB* (GEN 1C, 3A, 3C, 5B, 5C, 6C) had significantly better growth than the ancestor in 1 *µ*g/mL gentamicin (Fig. S1). Whilst the prevalence of *trkH* and *hyxB* was highly reproducible across the GEN populations as a whole, we did not find any instances of *trkH* and *hyxB* variants within the same sequenced strain (Table S1). Strikingly, none of the mutations identified in any of the strains from the GEN populations were found in any strains from the Glc+GEN populations. This suggests that the presence of glucose, at a concentration consistent with moderate glycosuria, alters the evolutionary response of UPEC to gentamicin.

**Fig. 3.**
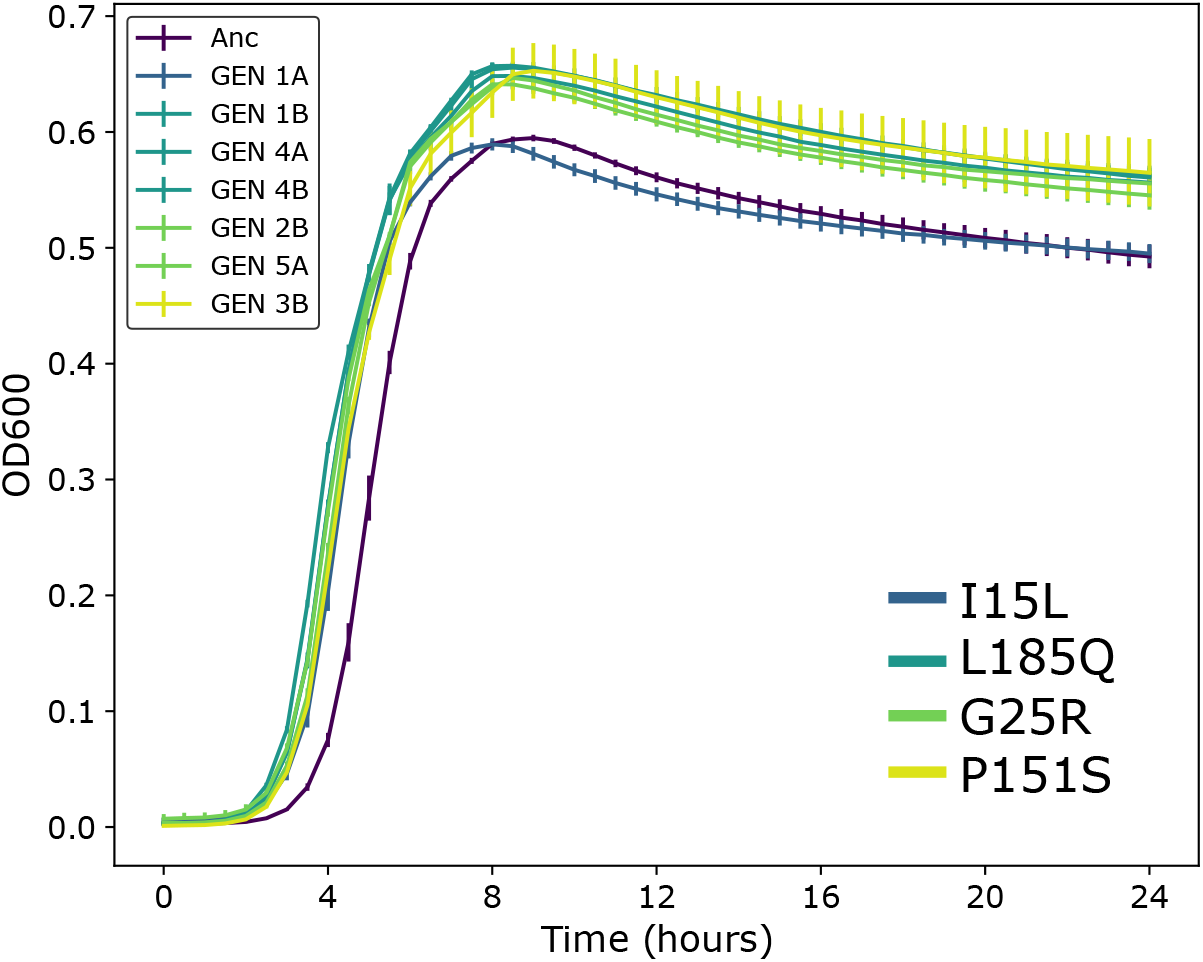
Growth kinetics in 1 *µ*g/mL gentamicin for the ancestor (Anc) and the strains with mutations in *trkH*. All curves are presented in Fig. S1 and area under the curve values in Fig. S2. Measurements in triplicate, error bars standard deviation.

## Discussion

Given the distinct metabolic microenvironment of the urinary tract and the extent to which is varies as a result of factors including diabetes mellitus and gestational diabetes, very little is known about how the concentration of different metabolites may drive UPEC evolution and response to antibiotics. Here, we discovered how a glucose concentration comparable to moderate glycosuria alters the response of a UPEC strain to gentamicin, an aminoglycoside prescribed in the UK for severe UTI or urosepsis (7; 8).

We identified several genes that were mutated in the presence of gentamicin only, including *trkH* and *hyxB*. The *trkH* gene encodes a potassium ion uptake system (30), and mutations in this gene have been observed in *E. coli* populations adapted to aminoglycosides, resulting in increased aminoglycoside resistance accompanied by increased susceptibility to nalidixic acid and tetracycline (31). The *hyxB* gene (also called, *upaB* (32)) encodes a virulence-associated autotransporter that mediates adherence to extracellular matrix proteins (32; 33; 34). It has been identified across *E. coli* pathotypes (33), with reports suggesting it is disrupted or absent in diarrheagenic *E. coli* but present in nosocomial, but not community, UPEC strains (35; 36; 37). The presence of both *trkH* and *hyxB* mutation in the same populations but, to the resolution of our sampling, never the same genome suggests that there may be selection for only one variant per strain.

Variants of the *rpoS* gene were identified in all twelve of the Glc and Glc+GEN populations, and not in either the control or the GEN populations. RpoS is a general stress sigma factor that is known to be induced during glucose starvation (38), with a role in osmotolerance (39). The lack of *rpoS* mutations in the GEN condition supports a recent study which found that expressing high levels of *rpoS* had minimal impact on antibiotic resistance (40). From our observations, it is reasonable to conclude that mutations in *rpoS* may be a result of a glucose-induced stress.

Taken together, our data suggest that a concentration of glucose consistent with moderate glycosuria may reduce the efficacy of antibiotics whilst also reducing the emergence of mutations linked to resistance. This may be due to increased efflux activity reducing the antibiotic concentration within the cell. AcrD is thought to efflux aminoglycosides in *E. coli* (41; 42), but analysis of a highly similar AcrD in *Salmonella* was found to not efflux gentamicin (43). More research is therefore needed to understand the mechanism underpinning these results, the effect of *trkH, hyxB*, and *rpoS* mutations on UPEC growth and antibiotic resistance, and the clinical ramifications for at-risk demographics.

## Acknowledgements

Whole genome sequencing was performed by MicrobesNG (http://www.microbesng.com). The research presented here was funded by a University of Birmingham College of Medicine and Health Research Development Fund award, awarded to RJH. RJH is supported by a BBSRC project grant (BB/W020602/1) awarded to AM. US-VAST002 is part of a strain collection donated to AM by James R Johnson. We are grateful to Jessica MA Blair for mechanistic insight.

## Data availability statement

The DNA dataset generated and analysed during the current study is available from NCBI BioProject with accession PRJNA1178604.

## Author contributions

SC: investigation. JS: investigation. AM: funding acquisition. RJH: conceptualization, methodology, formal analysis, investigation, data curation, writing - original draft, visualization, funding acquisition, supervision. All authors: writing - review & editing.

## Supplementary

**Fig. S1.**
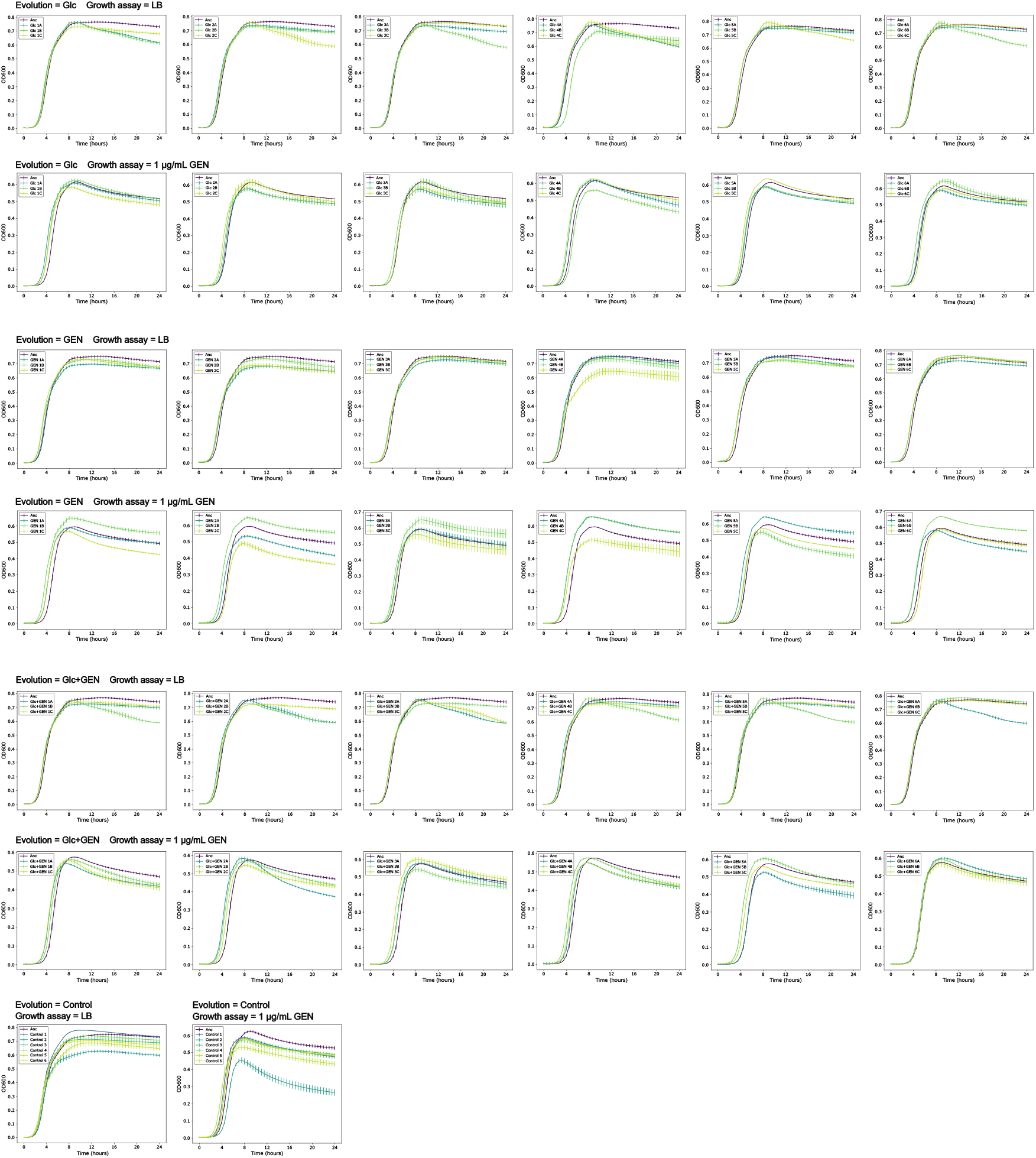
Growth kinetics of all sequenced strains in LB and in LB + 1 *µ*g/mL gentamicin. Plots are grouped by evolution condition (Glc, GEN, Glc+GEN, Control) and growth assay condition (LB, 1 *µ*g/mL GEN). Measurements in triplicate, error bars standard deviation.

**Fig. S2.**
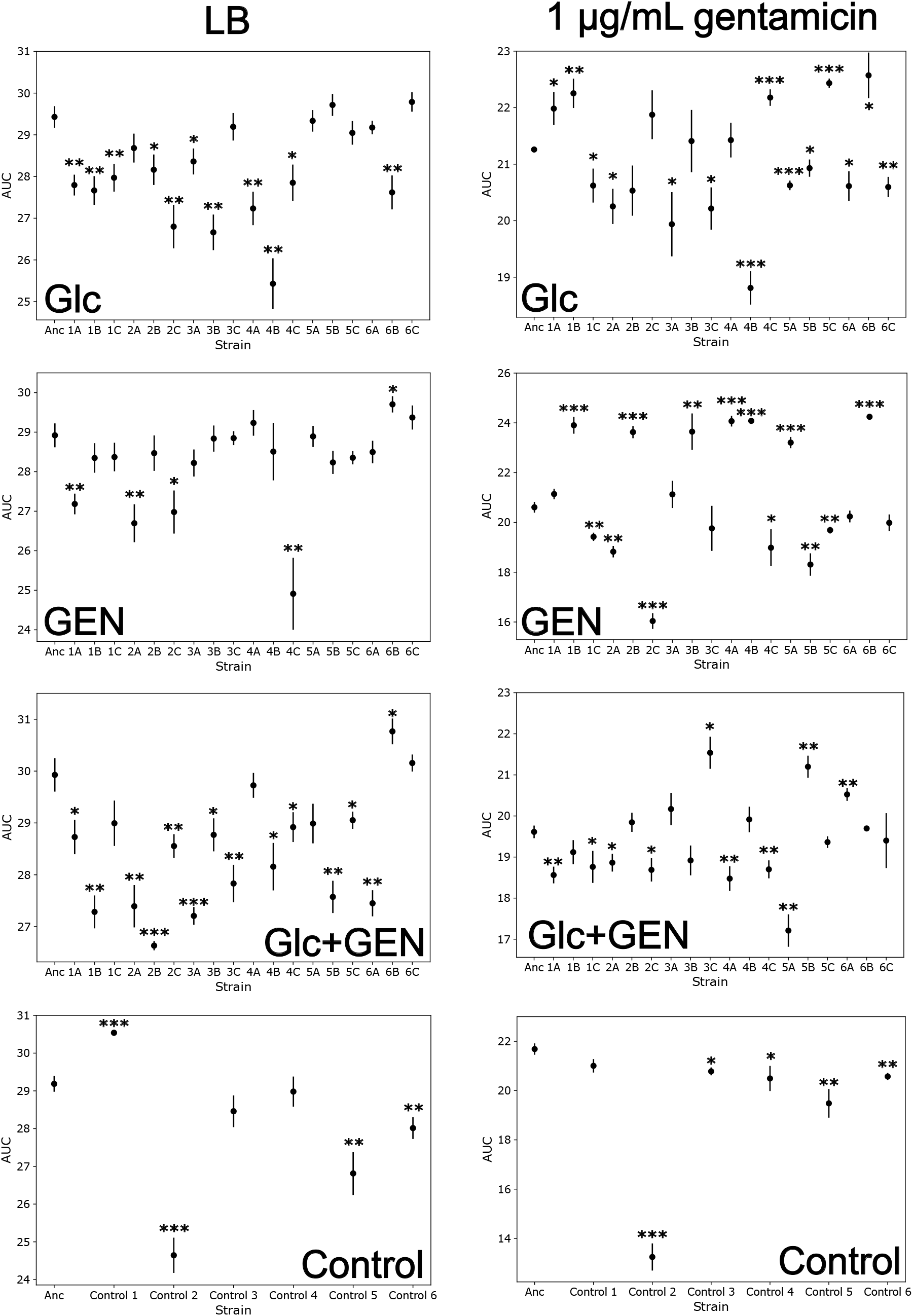
Area under the curve (AUC) values of growth kinetics in Fig. S1 in LB only (left) or 1 *µ*g/mL gentamicin. Measurements in triplicate, error bars standard deviation. Significant difference to the ancestral strain noted as * p < 0.05, ** p < 0.01, *** p < 0.001, one-way ANOVA.

